# Neuronal Modulation of Brown Adipose Activity Through Perturbation of White Adipocyte Lipogenesis

**DOI:** 10.1101/324160

**Authors:** Adilson Guilherme, David J Pedersen, Felipe Henriques, Alexander H. Bedard, Elizabeth Henchey, Mark Kelly, Kamal Rahmouni, Donald A. Morgan, Michael P Czech

## Abstract

White adipose tissue (WAT) secretes factors to communicate with other metabolic organs to maintain energy homeostasis. We previously reported that perturbation of adipocyte de novo lipogenesis (DNL) by deletion of fatty acid synthase (FASN) causes expansion of sympathetic neurons within white adipose tissue (WAT) and the appearance of “beige” adipocytes. Here we report evidence that white adipocyte DNL activity is also coupled to neuronal regulation and thermogenesis in brown adipose tissue (BAT). Induced deletion of FASN in all adipocytes in mature mice (iAdFASNKO) enhanced sympathetic innervation and neuronal activity as well as UCP1 expression in both WAT and BAT. In contrast, selective ablation of FASN in brown adipocytes of mice (iUCP1FASNKO) failed to modulate sympathetic innervation and the thermogenic program in BAT. Surprisingly, DNL in brown adipocytes was also dispensable in maintaining euthermia when UCP1FASNKO mice were cold-exposed. These results indicate that DNL in white adipocytes influences long distance signaling to BAT, which can modify BAT sympathetic innervation and expression of genes involved in thermogenesis.

## HIGHLIGHTS

- Inhibition of DNL in white adipocytes increases iWAT and BAT sympathetic neuron activities.
- Loss of FASN in white, but not brown adipocytes, enhances sympathetic innervation and thermogenesis in iWAT and BAT independent of thermoregulation.
- Suppression of DNL in iWAT appears to trigger BAT thermogenesis through a neural link.
- Although greatly increased in activity at 6°C, DNL in BAT is dispensable to maintain euthermia.

## INTRODUCTION

Adipose tissue is profoundly expanded in obesity based on greatly increased storage of neutral lipids. This expansion is frequently associated with the onset of metabolic diseases such as type 2 diabetes^1, 2, 3, 4^. Thus, understanding the functions and effects of adipose tissue on whole body metabolism is a major goal of this field. Animal studies have provided evidence that there are at least three different types of adipocytes, each presenting clear differences in their thermogenic potential and metabolic functions^5, 6, 7, 8^. White adipocytes, for example, display a unilocular lipid droplet with relatively low thermogenic potential and account for the increased stores of fatty acids in the form of triglyceride in obesity. Hydrolysis of this triglyceride in fasting conditions provides fatty acid fuel for other tissues^9, 10^. Brown adipocytes, on the other hand, contain multilocular lipid droplets, possess high thermogenic capacity and constitutively express the mitochondrial uncoupling protein 1 (UCP1). These cells utilize fatty acids and glucose as fuel to generate heat and maintain body temperature during cold-induced adaptive thermogenesis^5, 11, 12, 13^. White adipocytes can be converted into brown-like adipocytes, known as “brite” or “beige” adipocytes, which are also multilocular cells that express UCP1 and possess high thermogenic potential^5, 14^. This browning of white adipose tissue (WAT) is driven by release of catecholamines and perhaps other factors from sympathetic neurons within WAT that occurs during cold stimulus. Catecholamines signal through the cAMP pathway to stimulate lipolysis and upregulate UCP1, as well as other mitochondrial proteins that mediate fatty acid oxidation and the “beige” adipocyte phenotype^5, 14, 15^.

Conversely, multilocular brown adipocytes can be converted into white-like unilocular adipocytes with fewer mitochondria and lower oxidative capacity, UCP1 protein and thermogenic potential. This conversion of brown adipocytes into white-like adipocytes is known as whitening of brown adipose tissue (BAT) and occurs in situations such as at thermoneutral conditions (i.e., 30°C) where sympathetic innervation and activation within BAT is diminished. Altogether, these observations support the concept that under different physiological conditions, adipocytes are interconvertible cells that can adjust their appearance and metabolic phenotype as needed^7, 14, 16^. They are capable of changing from unilocular to multilocular, from anabolic to catabolic, from storing calories as lipids to dissipating calories as heat. These interconversions are distinct from differentiation of progenitor cells within adipose tissues which can also give rise to white, beige or brown adipocytes in response to sympathetic activity^14, 17^. Together, the combination of interconversion and differentiation mechanisms control the overall profile of adipocyte types within the tissue and these events are regulated by local sympathetic tone.

Recent work in our laboratory revealed that de novo fatty acid synthesis (DNL) within adipocytes may be linked to control of localized sympathetic nerve activity in adipose tissues ^3^. Thus, blockade of adipocyte DNL through selective inducible deletion of fatty acid synthase (*Fasn*) in adipocytes of mature mice enhanced sympathetic neuron innervation and browning of iWAT, along with an improvement of systemic glucose metabolism^3^. Therefore, adipocyte DNL perturbation not only modulates the thermogenic programming of iWAT, but also whole-body metabolism. Adipocyte DNL may produce bioactive lipids that alters adipose tissue functions^18^, energy balance and systemic metabolism^3, 19, 20^. This pathway is also a rich source of metabolites such as acetyl-CoA, malonyl-CoA, palmitate and lipid products, all known to control diverse cellular processes^2, 21, 22^. These metabolites mediate such post-translational protein modifications as protein acetylation^23^, malonylation^24^ and palmitoylation,^25^ which are implicated in histone regulation, gene expression and other cellular systems. Notably, the adipose DNL pathway is dynamically regulated by nutritional state, insulin and obesity^2, 3, 26, 27, 28^. In turn, these DNL perturbations may modulate adipose sympathetic activity. Taken together, the data available suggest the hypothesis that signaling by one or more small molecule metabolites connected to the DNL pathway within adipocytes mediates a signaling pathway that confers paracrine regulation of localized neurons.

Interestingly, while inducible deletion of FASN in both WAT and BAT (using adiponectin-Cre mice crossed to flox/flox FASN mice, denoted iAdFASNKO mice) in animals housed at 22°C caused expansion of sympathetic neurons and browning in WAT, no such effects were observed in BAT^3^. Furthermore, selective deletion of FASN in BAT (using UCP1-Cre mice crossed to flox/flox FASN mice, denoted UCP1-Cre-FASNKO) had no effect on either WAT or BAT in mice housed at 22°C. This absence of effect of FASN deletion in BAT was somewhat surprising since BAT is highly innervated and responsive to catecholamines. Also, DNL in BAT is vanishingly low at thermoneutrality but highly upregulated in cold-adapted mice. Based on these considerations, the present studies were designed to further investigate the role of DNL in BAT under these more extreme temperature conditions. Remarkably, deletion of FASN selectively in BAT in the UCP1-Cre-FASNKO mice did not decrease survival of mice at 6°C and had no detectable effect on UCP1 expression. Neither did selective FASN deletion in BAT have detectable effects on BAT function in mice housed at 30°C. However, deletion of FASN in both WAT and BAT in iAdFASNKO mice at 30°C did cause detectable sympathetic neuron expansion and increased UCP1 expression in BAT. These data indicate that DNL activity in white adipocytes within WAT can initiate long distance signaling to BAT that enhances its thermoregulation program.

## RESULTS

### Inducible deletion of adipocyte FASN stimulates sympathetic activity in adipose tissue

We first sought to establish whether adipocyte iAdFASNKO actually enhanced electrical activity of sympathetic neurons with adipose tissues in addition to the expansion of sympathetic innervation previously reported^3^. In these experiments, control and iAdFASNKO mice were first treated with tamoxifen (TAM) to induce FASN deletion. Four weeks later, the mice were anesthetized and the electrical activities of sympathetic fibers in iWAT and BAT were measured as previously described^29, 30^. Consistent with our previous results^3^, TAM treatment of iAdFASNKO mice, but not control mice, reduced the expression of FASN in adipocytes, increased the levels of adipose TH and UCP1 protein and induced the formation of UCP1-positive multilocular adipocytes within iWAT (Fig. 1A-C). Importantly, inducible knockout of adipose FASN enhanced the sympathetic nerve activity (SNA) from fibers innervating both iWAT and BAT (Fig. 1D-E and 1F-G). Altogether, these results demonstrate that suppression of DNL in adipocytes not only expands sympathetic innervation in iWAT, but also stimulates the activity of sympathetic fibers that innervate iWAT and BAT.

**Figure 1:**
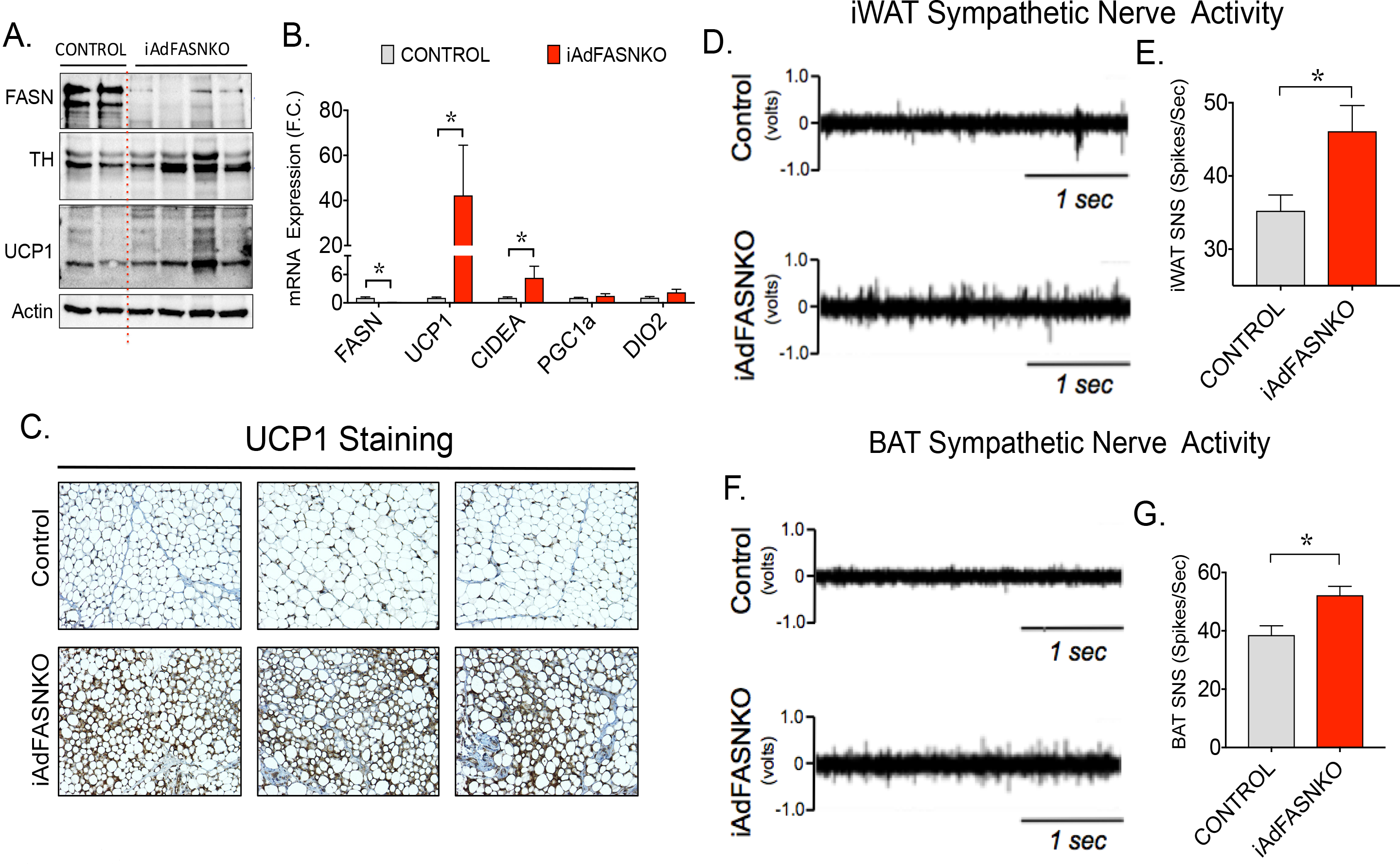
Inducible deletion of adipocyte FASN stimulates sympathetic and sensory nerve activities in adipose tissue. **(A**) Western blots of iWAT lysates from control and iAdFASNKO mice after TAM treatment. Indicated are FASN, TH, UCP1 and actin protein as loading control. **(B)** qRT-PCR was performed for quantifications of indicated gene expressions in iWAT from controls and iAdFASNKO mice. **(C)** Depicted is IHC analysis for detection of UCP-1 protein in iWAT from control, iAdFASNKO. Results are representative image of 8-12 mice per group. **(D-G)** Representative sympathetic nerve activity recordings and respective averages in iWAT and in BAT. Graphs show the mean +/− SEM. N = 12 mice per group; \**p* < 0.05.

### De novo lipogenesis in BAT is not required to maintain euthermia

Sympathetic neuron modulation in iWAT and BAT elicited by adipocyte FASN deletion shown in Fig. 1 suggests that DNL metabolites in adipocytes might influence adipose sympathetic drive needed during cold-induced adaptive thermogenesis. Indeed, a number of studies reported that DNL gene expression is strongly upregulated in BAT in cold-challenged mice^28, 31^, and it has been suggested that the DNL pathway may also be required for providing optimal fatty acid fuels to sustain cold-induced thermogenesis in BAT and euthermia ^28, 32^. To assess whether DNL in brown adipocytes is necessary for BAT thermogenesis and proper control of body temperature during cold-exposure, UCP1-Cre-FASNKO mice and FASN^flox/flox^ littermates that do not express Cre-recombinase were subjected to an ambient temperature of 6°C. As shown in Fig. 2, specific deletion of FASN in BAT, but not in iWAT, from the UCP1-Cre-FASNKO mice was confirmed. Importantly, selectively deleting FASN in brown adipocytes did not affect cold-induced UCP1 expression in BAT or browning of iWAT (Fig. 2A and 2F). Consistently, UCP1-Cre-FASNKO mice have normal body temperature and do not become hypothermic when acutely challenged with cold temperature (6°C) (Fig 2B). Moreover, deletion of FASN in brown adipocytes of cold-challenged mice did not affect the body weight or BAT mass and morphology (Fig. 2C-E), nor did it affect induction of UCP1 protein in iWAT of these cold-challenged mice lacking FASN in BAT (Fig. 2F). Additionally, iWAT mass is unchanged in cold adapted UCP1-Cre-FASNKO mice (Fig. 2G). Taken together, these results indicate that DNL in brown adipocytes is not necessary for normal thermogenesis in BAT and for proper control of body temperature in mice
exposed to 6°C conditions.

**Figure 2:**
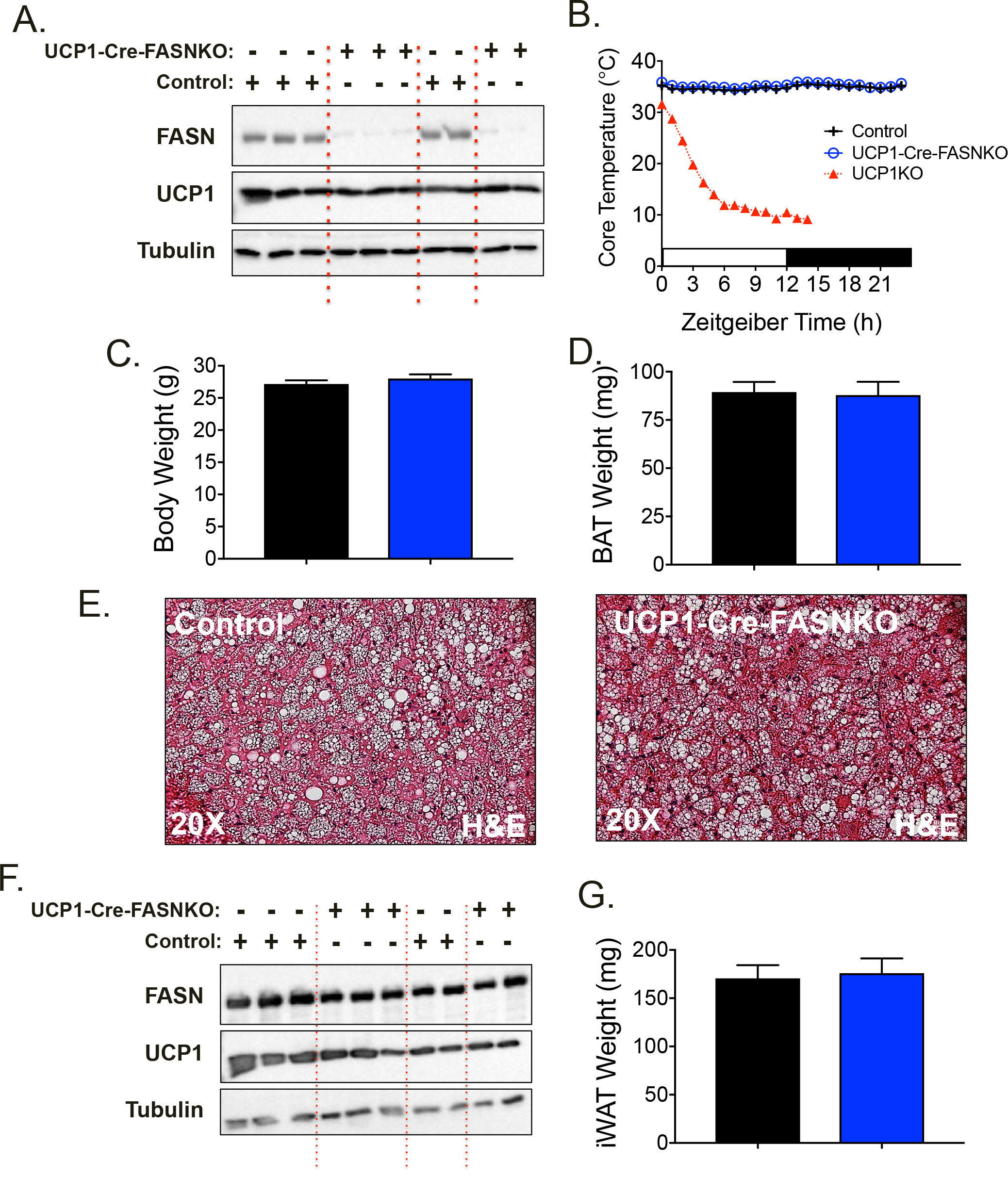
Fatty acid biosynthesis in brown adipocytes is not required to maintain euthermia. **(A)** Western blots of BAT lysates from control and UCP1-Cre-FASNKO mice after 1 week at 6°C. Indicated are FASN, UCP1 and tubulin protein as loading control. **(B)** Core temperature in acute cold challenge (6°C), starting from room temperature in control, UCP1-FASNKO and UCP1-knockout mice. **(C)** Total body weight and (**D**) brown adipose tissue depot weight after cold exposed at 6°C for 1 week. **(E)** Representative H&E images of BAT from control and UCP1-FASNKO mice cold-exposed at 6°C for 1 week. **(F)** Western blots of iWAT lysates from control and UCP1-FASNKO mice cold exposed at 6°C for 1 week. **(G)** Inguinal adipose tissue mass from control and UCP1-FASNKO cold 30 exposed at 6°C for 1 week. Graphs show the mean +/− SEM. N= 5-7 mice per group.

We next conducted experiments in iAdFASNKO mice to investigate further whether inhibition of DNL in all mouse adipocyte types would affect cold-induced thermogenesis required to control body temperature. iAdFASNKO mice were acutely cold exposed using the same protocol as in Fig. 2B. Similar to the UCP1-Cre-FASNKO mice, iAdFASNKO mice have normal body temperature, body weight, BAT and iWAT mass, and do not become hypothermic when challenged with cold temperature (Supplementary Fig. 1). Taken together with the results from UCP1-Cre-FASNKO mice, we conclude that DNL in all adipocytes types is dispensable for cold-induced thermogenesis required to maintain euthermia.

### Adipocyte FASN KO expands adipose sympathetic neurons even at thermoneutrality

A key potential caveat in our previous report^3^ and in Fig 1 is that enhanced adipose innervation and browning in iAdFASNKO mice at 22°C is simply due to increased heat loss in the animal, due to decreased dermal or tail insulation^11, 12^. We therefore investigated whether the enhanced SNS expansion and browning of adipose tissue depots could also be detected in iAdFASNKO mice housed at thermoneutrality (TN, 30°C). After three weeks of TN housing, control and iAdFASNKO mice were treated with TAM to induce FASN deletion in adipocytes in the floxed mice. Two weeks following TAM treatment, the mice were treated with the β-3 agonist CL316,243 or PBS (1 i.p. injection daily for 6 days) to assess the effects of Adrb3 activation on the thermogenic program of BAT and WAT at TN. As shown in Fig. 3A-B, adipocyte FASN knockout at TN caused increased TH signal in immunohistochemistry (IHC) analysis and induced the appearance of multilocular adipocytes within the iWAT, mimicking the effect of β-3 agonist, although to a lesser degree (Fig. 3D). Importantly, at 30°C, UCP1 expression in iWAT was abolished, an important positive control showing that thermoneutral conditions were achieved (Fig. 3C and 3F). Interestingly, little to no effect from CL316,243 treatment or from FASN knockout was observed on UCP1 expression at this thermoneutral condition (Fig. 3C and 3F). Although the combination of adipocyte FASN deletion plus CL316,243 treatment appears to be additive in enhancing multilocularity of adipocytes and UCP1 expression at TN, these effects were modest when compared to the browning detected in iAdFASNKO mice at 22°C (Fig. 3C-3F). Altogether, these data indicate that loss of FASN in mouse adipocytes causes significant increases in TH expression, SNS expansion and multilocular adipocyte appearance in iWAT, even at thermoneutral temperature, and therefore is not dependent on thermoregulation for inducing these effects.

**Figure 3:**
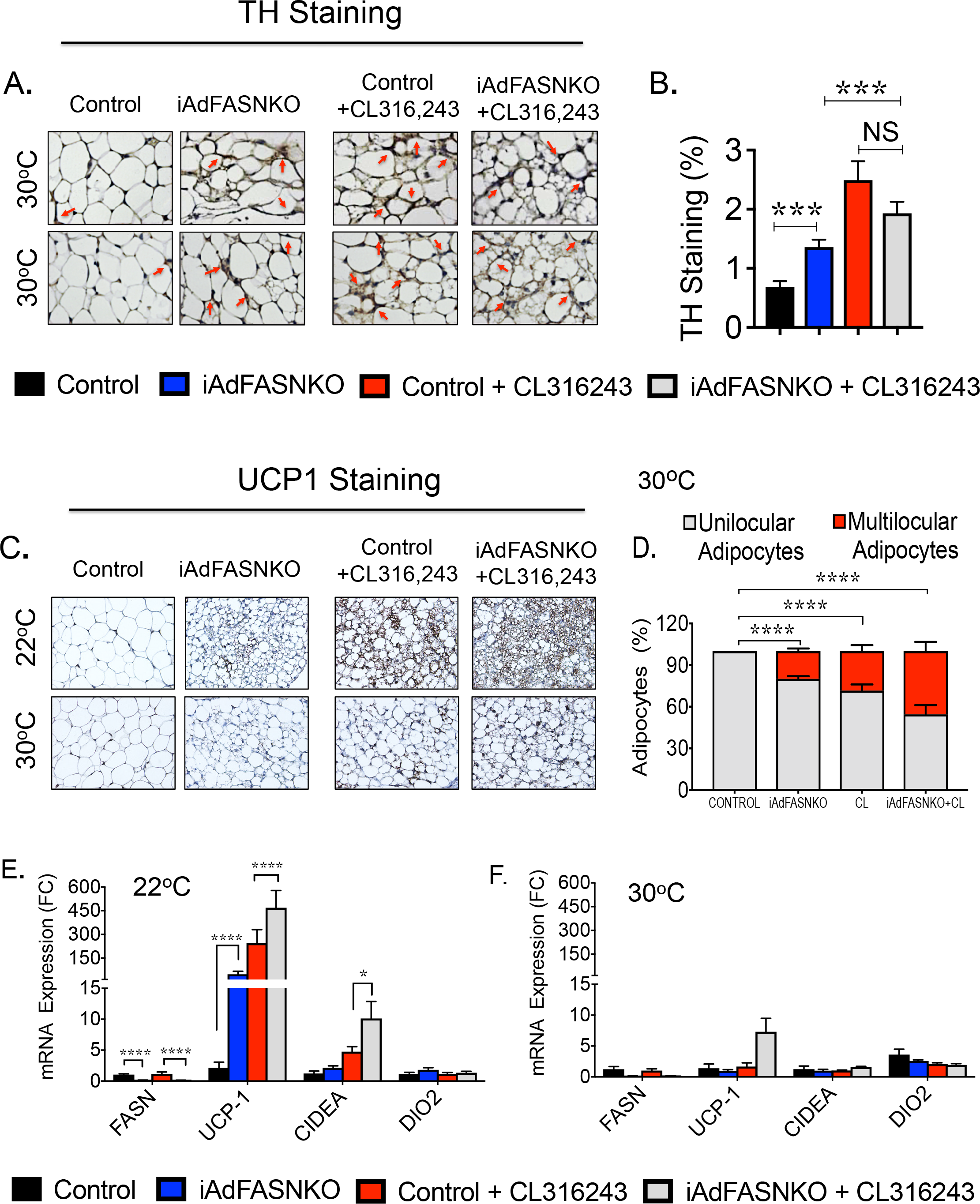
Suppression of fatty acid biosynthesis in adipocytes enhances SNS expansion and browning in iWAT, even at thermoneutrality. **(A)** Immunohistochemistry (IHC) for detection of Tyrosine hydroxylase (TH) contents in iWAT from control iAdFANSKO mice housed at thermoneutrality (30°C), treated or not with CL316,243 for 6 days. Loss of FASN in adipocytes increased TH levels in iWAT and CL316,243 treatment further enhanced TH contents in iWAT. **(B)** Quantification of TH staining in iWAT from images depicted in **(A). (C)** Depicted are IHC analyses for detection of UCP-1 protein in iWAT from control, iAdFASNKO mice housed at thermoneutrality and treated or not with CL316,243. Multilocular adipocytes are detected in iWAT from iAdFASNKO mice, while detection of UCP1 protein requires adipose FASN deletion plus CL316,243 treatment. **(D)** Quantification of multilocular adipocytes and UCP1 staining in iWAT from images depicted in **(C). (E-F).** qRT-PCR was performed for quantifications of indicated genes in iWAT from controls and iAdFASNKO mice housed either at 22°C **(E)** or at 30°C **(F)** and treated or not with CL316,243 for 6 days. Graphs show the mean +/− SEM. N= 5-7 mice per group. * P < 0.05; **** P < 0.0001.

### Adipocyte FASN deficiency enhances innervation and attenuates whitening of BAT at thermoneutrality

We next conducted experiments to examine whether the increased SNS expansion and appearance of multilocular adipocytes observed in iWAT from iAdFASNKO mice housed at TN conditions could also be detected in BAT. As depicted in Fig. 4A, BAT from mice kept at 30°C for 6 weeks loses its multilocular appearance, and TH-positive neuron content was also found to be strongly reduced by TN. Consistent with the whitening of BAT that occurs under TN conditions^16, 33^, BAT UCP1 expression levels were also markedly suppressed at 30°C (Fig. 4E-G). Interestingly, similar to the effect noted in iWAT (Fig. 3A-B), deletion of FASN in all adipocytes increased TH-positive neuron content and promoted the formation of multilocular brown adipocytes in BAT at TN (Fig. 4A-D). As expected, CL316,243 treatment partially restored both the adipocyte multilocularity and sympathetic neuron density in BAT from control mice, while treatment of iAdFASNKO mice with this β3-agonist completely restored the sympathetic innervation and multilocular appearance of BAT, despite the mice being housed at TN (Fig. 4A-D). Immunoblotting to detect TH protein levels in tissue lysates confirmed the increase in TH-positive neurons in BAT from iAdFASNKO mice at TN treated with CL316,243 (Fig. 4C-D).

**Figure 4:**
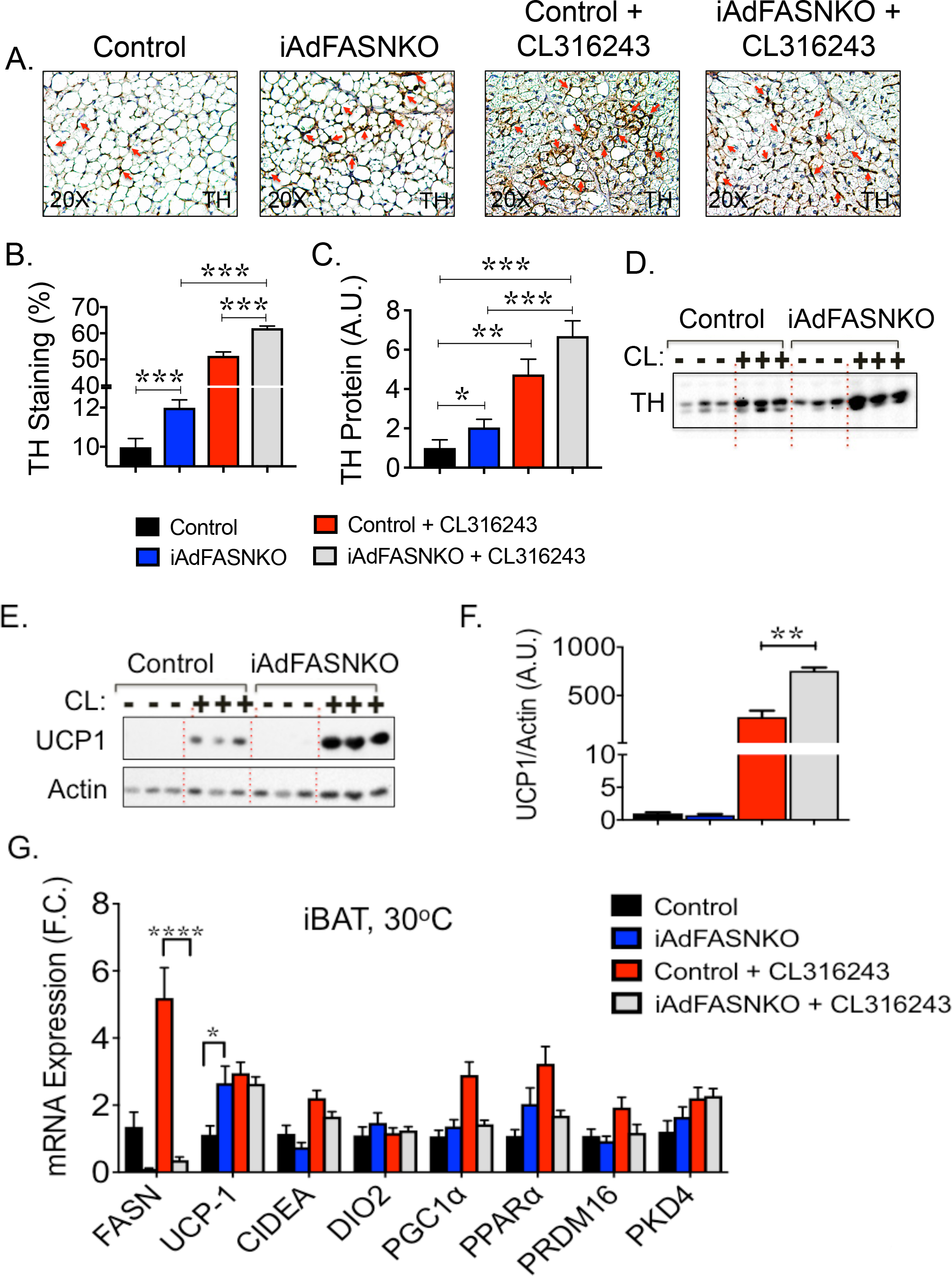
Inhibition of fatty acid biosynthesis in adipocytes enhances SNS expansion and thermogenic protein in BAT from mice housed at thermoneutrality. **(A)** Immunohistochemistry (IHC) for detection of Tyrosine hydroxylase (TH) contents in BAT from control iAdFANSKO mice housed at thermoneutrality (30°C), treated or not with CL316,243 for 6 days. Depletion of FASN in adipocytes increased TH levels in BAT and CL316,243 treatment further enhanced TH contents in BAT. Red arrows indicate TH-positive neurons. **(B)** Quantification of TH staining in BAT from images depicted in **(A). (C)** Quantification of TH protein levels in BAT upon induction of adipocyte FASN deletion. TH protein levels were quantified by densitometry from immunoblot data shown in **(D).** In **(D)** Westerns blot for detection of TH protein in BAT from control and iAdFASNKO mice housed at thermoneutrality and treated or not with CL316,243. **(E)** Westerns blot for detection of UCP1 protein in BAT from control, iAdFASNKO mice housed at thermoneutrality and treated or not with CL316,243. **(F)** Quantification of UCP1 protein levels in BAT upon induction of adipocyte FASN deletion. UCP1 protein levels were quantified by densitometry from immunoblot data shown in **(E). (G).** qPCR was performed for quantifications of indicated genes in iWAT from controls and iAdFASNKO mice housed either at 30°C and treated or not with CL316,243 for 6 days. Graphs show the mean +/−SEM. N= 5-7 mice per group. * P < 0.05; ** P < 0.001; **** P < 0.0001.

Interestingly, while deletion of FASN in adipocytes seems to be sufficient to partially restore sympathetic innervation in BAT and to increase the appearance of multilocular adipocytes (Fig. 4A-B) at TN, the enhanced SNS expansion was not accompanied by increased UCP1 expression, as depicted in Fig. 4E-F. However, while CL316,243 treatment alone at TN partially upregulated UCP1 expression, the combination of adipocyte FASN deletion plus CL316,243 treatment completely restored the UCP1 expression levels in BAT. Overall, these results are consistent with the hypothesis that inhibition of adipocyte DNL through FASN knockout in both white and brown adipocytes activates signaling pathways that promotes sympathetic innervation and formation of multilocular adipocytes in both iWAT and BAT, even at TN conditions, thus attenuating whitening of BAT at TN.

### Brown adipocyte FASN deficiency fails to expand SNS and attenuate whitening of BAT at TN

Our previous studies demonstrated that while deletion of FASN in both white and brown adipocytes promotes sympathetic innervation and browning in iWAT from mouse housed at 22°C, selective deletion of FASN in only brown adipocytes does not affect adipose TH content or browning and UCP1 expression in iWAT or BAT^3^. However, whether selective inhibition of DNL in brown adipocytes in mice housed in TN conditions could cause “browning” of the “whitened” BAT, similar to the phenotype seen with deletion of FASN in both white and brown adipocytes in iAdFASNKO mice (Fig. 3–4), was not previously addressed. To do this, control (FASN-flox/flox) and iUCP1-Cre-FASNKO mice were housed at TN for 3 weeks and then treated with TAM to induce FASN deletion specifically in brown adipocytes in the latter mice. Three weeks following TAM induction, mice were treated with CL316,246 or PBS (1 i.p. injection daily for 6 days). Adipose tissue was then harvested and processed to assess the effects of selective brown adipocyte FASN deletion on TH-positive neuron content, UCP1 and thermogenic gene expression levels in adipose tissue depots.

Consistent with the results shown in Fig. 4C-D, treatment of control mice with CL316,243 partially restored the TH and UCP1 expression levels in BAT at TN conditions (Fig. 5A-B). However, selective deletion of FASN in brown adipocytes failed to increase TH and UCP1 expression levels in BAT from mice kept at TN. Importantly, loss of FASN in only brown adipocytes also failed to further enhance the TH contents (Fig. 5D), UCP1 protein and expression of several thermogenic genes in BAT from mice treated with CL316,243 at TN (Fig. 5A-E). Taken together, these results indicate that while deletion of FASN in white and brown adipocytes initiates signals to expand and activate local neurons and induce adipose browning, selective inactivation of FASN in brown adipocytes is not sufficient to trigger such signals. Therefore, we hypothesize that the DNL pathway in white adipocytes produces intermediate metabolites that initiate signaling pathways to local neurons that then mediate effects in distant BAT (Fig. 5F).

**Figure 5:**
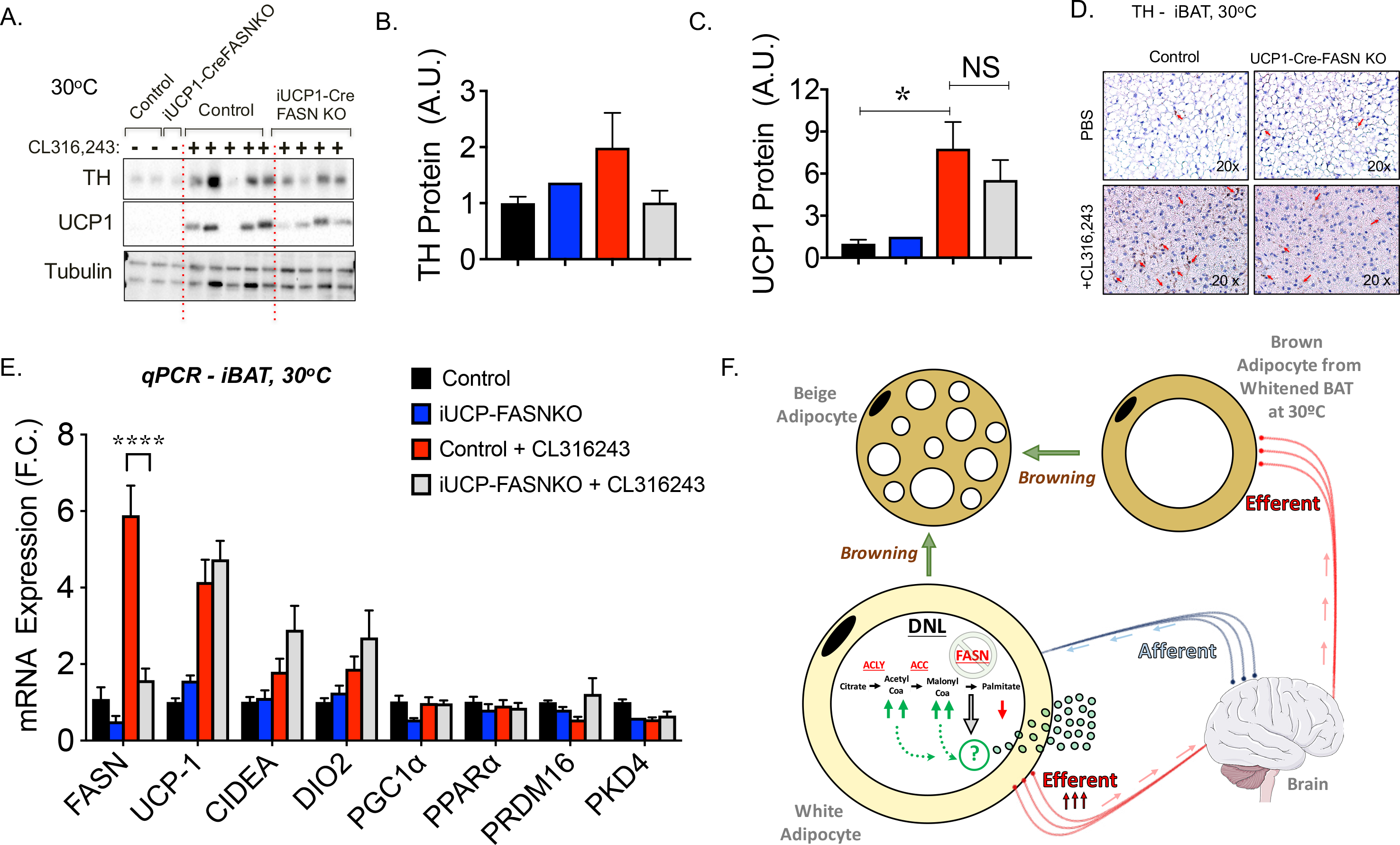
Suppression of fatty acid biosynthesis selectively in brown adipocytes does not affect SNS expansion in BAT or extent of multilocular cells in iWAT at thermoneutrality. **(A)** Westerns blot for detection of TH, UCP1 and tubulin protein in BAT from control, iUCP1-Cre-FASNKO mice housed at thermoneutrality and treated or not with CL316,243. **(B)** Quantifications of TH (**B**) and UCP1 **(C)** protein levels in BAT upon induction of adipocyte FASN deletion. TH protein levels were quantified by densitometry from immunoblot data shown in **(A). (D)** Immunohistochemistry (IHC) for detection of Tyrosine hydroxylase (TH) contents in BAT from control mice or UCP1-Cre-FANSKO mice housed at thermoneutrality. **(E)** q-PCR was performed for quantifications of indicated genes in iWAT from controls and iUCP1-Cre-FASNKO mice housed at 30°C and treated or not with CL316,243 for 6 days. **(F)** Proposed model show how disruption of adipocyte DNL pathway through FASN knockout stimulate adipose neuronal activity and thermogenic program. Accordingly, adipocyte FASN deletion enhances the levels of intermediate lipids acetyl-CoA and malonyl-CoA (green arrows), but reduces palmitate (red arrows). Such changes trigger the production of neurotropic factors and signals conveyed to brain. Graphs show the mean +/−SEM. N= 5-7 mice per group. * P < 0.05; **** P < 0.0001.

## DISCUSSION

The major finding in the present study shows that deletion of FASN in all mature white and brown adipocytes in iAdFASNKO mice enhances local sympathetic innervation of iWAT and BAT even at TN, while FASN deletion in only brown adipocytes in UCP1-Cre-FASNKO mice has no such effect (Fig. 3–5). BAT also displays increased expression of UCP1 mRNA in TAM treated iAdFASNKO mice at TN, but not in UCP1-Cre-FASNKO mice. There was a similar failure of the UCP1-Cre-FASNKO mouse to show additive effects of CL316,243 and FASNKO on UCP1 and TH expression in BAT, as is observed in the iAdFASNKO mice (Fig. 4 and 5). Also in line with these divergent results in the two FASN KO mouse models, our previous studies showed no effect of brown adipocyte FASN deletion on glucose tolerance, whereas iAdFASNKO mice displayed clear improvement in glucose tolerance^3^. Both the iAdFASNKO and UCP1-Cre-FASNKO mice at TN do show the expected decrease in FASN expression in either iWAT and BAT or only BAT, respectively, indicating that enough *Ucp1* promoter activity is available to mediate Cre expression in the UCP1-Cre-FASNKO mice at TN. These data strongly suggest that FASN KO only in white adipocytes can mediate the effects of increased TH and UCP1 expression on BAT. In line with the above conclusions, induced FASN deletion in adipocytes in mature iAdFASNKO mice also enhanced both WAT and BAT SNS activity.

The most likely route of such communication between WAT and BAT is through sensory neurons in WAT that direct the CNS to enhance sympathetic activity in BAT. This interpretation is supported by the data in Fig. 1 showing sympathetic nerve activity in BAT is increased upon FASN deletion in the iAdFASNKO mice. Furthermore, the results are consistent with a working model whereby suppression of DNL in white adipocytes initiates regulatory signals that promote not only increased sympathetic innervation density in iWAT and BAT, but also increased activity of the expanded nerve fibers (Fig. 1). Other studies have demonstrated increased activity of sympathetic nerves in iWAT and BAT after cold stimulus^34, 35^ and upon central nervous system activation such as direct administration of FGF21 peptide in the brain^36^ or via local optogenetic activation of sympathetic fibers ^37^. The idea that communication occurs between WAT and BAT in both directions has also been previously described. For example, directed activation of iWAT lipolysis was found to activate local iWAT afferents triggering a neural circuit from WAT to BAT that induces BAT thermogenesis^38, 39^. In addition, a number of studies utilizing adipose tissue denervation procedures have demonstrated neuronal communication between different white adipose fat depots^40, 41, 42^. The present study extends these observations by showing that perturbations in adipocyte fatty acid metabolism can also have striking effects on local and distant sympathetic nerve activities, along with browning of iWAT and BAT. These results implicate that lipid intermediates and/or fatty acids from the DNL pathway in WAT control local afferent nerves and therefore a neuronal circuit that regulates iWAT browning and systemic metabolism, as illustrated in Figure 5F. According to this model afferent signals originated in white adipocytes deficient in FASN are conveyed to the CNS to drive sympathetic outflow in adipose tissue promoting browning (Fig. 5F).

One mechanism whereby adipocyte FASN knockout might enhance the SNS is through promoting heat loss in mice. As fatty acid biosynthesis in iWAT might be necessary for optimal body insulation, it is conceivable that inactivation of FASN in adipocytes could disrupt proper skin insulation, triggering thermoregulatory reflexes^12^. It is also conceivable that cold perception could be heightened in iAdFASNKO mice, thereby necessitating increases in sympathetic innervation, iWAT browning and basal thermogenesis. Accordingly, a similar mechanism has been proposed to explain how the deficiency of the lipogenic enzyme stearoyl-CoA desaturase-1 enhances thermogenesis in mice^43^. However, this does not appear to be the case in the present iAdFASNKO mouse model since increased TH-positive neurons in both iWAT and BAT are still observed when these mice are caged at TN (Fig. 3 and 4). However, UCP1 induction in iWAT was not observed in response to adipocyte FASN knockout under TN conditions (Fig. 3). Thus, our findings suggest that while adipocyte FASN knockout increases SNS innervation and multilocular cells in adipose tissue, the induction of UCP1 and the complete thermogenic program does require mildly cold temperatures such as 22°C and likely CNS participation. Based on these overall results, we conclude that the effect of adipocyte FASN deletion on local sympathetic nerve density and browning is not due to a compensatory mechanism in order to cope with systemic heat loss.

Our study also revealed unexpected results related to the contribution of DNL to cold-induced thermogenesis in BAT and the ability of mice to maintain euthermia in the cold. It has long been known that cold exposure in mice greatly increases the expression of DNL enzymes and activity in BAT^31, 44^, and more recently studies showed the DNL pathway positively correlates with thermogenesis in BAT^28, 32^. Moreover, it has been proposed that fatty acids are required to drive thermogenesis by directly activating UCP1^45^ and by providing fuel for high rates of oxidative metabolism^13^. Altogether, these observations support the concept that increasing DNL in BAT would likely be essential for survival at very low temperatures, as it would contribute to the fatty acid supply for oxidation during thermogenesis. However, to our knowledge, the requirement of DNL for thermogenesis in BAT and maintenance of euthermia during cold exposure has not been formally tested. Surprisingly, we did not detect significant differences in expression of UCP1 protein or other thermogenic genes in BAT or in body temperature of UCP1-Cre-FASNKO mice devoid of brown adipocyte FASN when compared to control mice (Fig. 2). These results indicate that DNL in BAT is dispensable for cold-induced thermogenesis and maintenance of euthermia in cold exposed mice.

In summary, we show here that suppression of FASN in white adipocytes of iAdFASNKO mice at room temperature elicits a remarkable neuromodulation of sympathetic nerves, both by expansion and increased electrical activity, in WAT and BAT. This adipocyte-neuron crosstalk may be essential for adipose inter-tissue communication, regulation of adipose thermogenesis and systemic metabolic homeostasis. Mechanistically, how inactivation of FASN in white adipocytes promotes such strong neuromodulation of nerve fibers in adipose tissue at the molecular level remains unknown. We are currently performing experiments to address this important question.

## METHODS

### Animal studies

Mice were housed on a 12 h light/dark schedule and had free access to water and food, except when indicated. Mice with conditional FASN^flox/flox^ alleles were generated as previously described^18^. To selectively delete FASN in adipocytes from adult mice, homozygous FASN^flox/flox^ animals were crossed to Adiponectin-Cre-ERT2 mice to generate the TAM-inducible, adipocyte-specific FASN knockout mice referred to as iAdFASNKO (Guilherme et al., 2017). At eight-weeks of age, both control FASN^flox/flox^ and iAdFASNKO were treated via intraperitoneal (i.p.) injection once a day with 1 mg TAM dissolved in corn oil for 6 consecutive days. FASN^flox/flox^ animals were also crossed with iUCP1-Cre-ERT2 mice (Jackson Laboratory) to generate the iUCP1-Cre-FASNKO that specifically delete FASN in brown adipocytes upon TAM treatment as described. The UCP1 KO mice were obtained from JAX Laboratory (Jackson Laboratory stock number 017476).

### Mice housing at TN and CL316,243 treatment

For the effects of thermoneutrality on adipose tissue innervation and browning, 8-week-old control mice and iAdFASNKO mice were transferred from 22°C (mild-cold) to 30°C (thermoneutrality) and acclimated for 3 weeks. Then, control and iAdFASNKO mice were treated with tamoxifen (TAM) as previously described (Guilherme et al., 2017). Two weeks post-TAM, mice were treated with 20 mg/kg of β3-agonist CL316,243 or PBS (one daily i.p. injection for 6 days). On the next day, adipose tissue was harvested and processed for histological and biochemical analyzes.

### Cold Challenge

To assess cold tolerance, control and knockout mice were placed at 6 °C in the morning and provided free access to food and water. Rectal temperatures were recorded every 1 h for a total of 24h.

### Sympathetic and Sensory Nerve Recordings

For determination of sympathetic and sensory nerve activities in adipose tissue from iAdFASNKO mice, the procedure was performed as follows: Each mouse was anesthetized with intraperitoneal administration of ketamine (91 mg/kg body weight) and xylazine (9.1 mg/kg body weight). Tracheotomy was performed by using PE-50 tubing to provide an unimpeded airway for the mouse to breathe O_2_-enriched room air. Next, a micro-renathane tubing (MRE-40, Braintree Scientific) was inserted into the right jugular vein for infusion of the sustaining anesthetic agent: α-chloralose (initial dose: 12 mg/kg, then sustaining dose of 6 mg/kg/h). A second MRE-40 catheter inserted into the left common carotid artery was connected to a Powerlab via a pressure transducer (BP-100; iWorx Systems, Inc.) for continuous measurement of arterial pressure and heart rate. Core body temperature was monitored through a rectal probe and maintained at 37.5° C. Next, each mouse underwent direct multifiber recording of sympathetic nerve activity (SNA) from a nerve innervating white adipose tissue (WAT) followed by SNA subserving brown adipose tissue (BAT). The nerve subserving the inguinal WAT was accessed through a small incision made on the right flank near the hindlimb. The sympathetic nerve fascicle was carefully isolated from surrounding connective tissues. A bipolar platinum-iridium electrode (40-gauge, A-M Systems) was suspended under the nerve and secured with silicone gel (Kwik-Cast, WPI). The electrode was attached to a high-impedance probe (HIP-511, Grass Instruments) and the nerve signal was filtered at a 100- and 1000-Hz cutoff with a Grass P5 AC pre-amplifier and amplified 10^5^ times. The filtered and amplified nerve signal was routed to a speaker system and to an oscilloscope (model 54501A, Hewlett-Packard) to monitor the audio and visual quality of the SNA recording. The nerve signal was also directed to a resetting voltage integrator (model B600c, University of Iowa Bioengineering) and finally to a MacLab analog-digital converter (Model 8S, AD Instruments Castle Hill, New South Wales, Australia) containing software (MacLab Chart Pro; Version 7.0) that utilizes a cursor to analyze the total activity and to count the number of spikes/second that exceeds the background noise threshold.

After achieving a stable anesthetic and iso-thermal condition, a continuous recording of baseline inguinal SNA was measured over a 30 min control period. The exposed central end of the inguinal nerve fiber was then carefully cut to record the afferent inguinal SNA for 30 minutes. At the conclusion, the afferent ends of the sympathetic nerve fiber were cut and the residual background noise used to normalize the intact and afferent basal inguinal SNA.

After completing the WAT SNA recording, the mouse was repositioned to gain access to the nerve fascicle that innervates the inter-scapular BAT through an incision in the nape of the neck. The BAT sympathetic nerve was isolated and instrumented for SNA recording as above. Once the anesthetic and iso-thermal conditions are stable, continuous recording of baseline BAT SNA was measured over a 30 min control period. At the end of this period, the exposed central end of the BAT sympathetic nerve fiber was carefully cut, and the recording of the afferent BAT SNA was performed for 30 minutes. At the conclusion of the study, the distal end of the BAT sympathetic nerve fiber was severed, and the residual background noise used to normalize the intact and afferent basal BAT SNA.

### Westerns Blots

For protein expression analyses, adipose tissues from indicated mice were homogenized in lysis buffer (20 mM HEPES [pH 7.4], 150 mM NaCl, 2 mM EDTA, 1% Triton X-100, 0.1% SDS, 10% glycerol, 0.5% sodium deoxycholate) that had been supplemented with Halt protease and phosphatase inhibitors (Thermo Pierce). Samples from tissue lysates where then resolved by SDS-PAGE and immunoblots were performed using standard protocols. Membranes were blotted with the following antibodies: FASN (BD Biosciences), TH (Abcam and Millipore), UCP1 (Alpha Diagnostics), Actin, Tubulin (Sigma-Aldrich).

### Histological Analysis

For the immunohistochemistry (IHC), tissue samples were fixed in 4% paraformaldehyde and embedded in paraffin. Sectioned slides were then stained for UCP1 (Abcam) and TH (Abcam and Millipore) at the UMASS Medical School Morphology Core. UCP1- and TH-immunostained sections from WAT were used to measure the percentage of positive staining per group. For IHC quantification random areas of adipose tissues from control and iAdFASNKO mice were selected and quantified by IHC - Image Analysis Toolbox plugin by ImageJ software. The quantification represents the average for six fields of five animals per group observed under final magnification of 400x. Scale bar = 50 μm. The morphometry of adipocytes was performed in serial paraffin sections of WAT stained with hematoxylin and eosin (H&E). For this study, we used six different fields of five animals per group. Counts were performed to determine the number of unilocular and multilocular cells in the adipose tissue. We obtained the specific number of these different types of cells (unilocular and multilocular adipocytes) and divided by total number of cells to obtain the total percentage. The images of the unilocular and multilocular adipocytes were acquired by a digital light microscope at 200× final magnification. The areas were profiled manually using ImageJ software. Scale bar = 100 μm.

### RNA isolation and RT-qPCR

Total RNA was isolated from mouse tissue using QIAzol Lysis Reagent Protocol (QIAGEN) following the manufacturer’s instructions. cDNA was synthesized from 1 ◻Lysis Reagent ProtocoiScript cDNA Synthesis Kit (BioRad). Quantitative RT-PCR was performed using iQ SybrGreen supermix on a BioRad CFX97 RT-PCR system and analyzed as previously described^46, 47^. *36B4*, *Hprt and Gapdh* served as controls for normalization. Primer sequences used for qRT–PCR analyses were listed in Supplementary Table 1.

### Statistical analysis

Data were analyzed in GraphPad Prism 7 (GraphPad Software, Inc.). The statistical significance of the differences in the means of experimental groups was determined by Student’s *t*-test as indicated. The data are presented as means ± SE. P values ≤ 0.05 were considered significant.

## ACKNOWLEDGMENTS

We thank all members of Michael Czech’s Lab for helpful discussions and critical reading of the manuscript. We thank the UMASS Morphology Core for assistance in immunohistochemistry analysis, and Clay Semenkovich and Irfan Lodhi for the generous gift of FASN ^flox/flox^ mice. This work was supported by NIH grants DK30898 and DK103047 to M.P.C.

## AUTHOR CONTRIBUTIONS

A.G., D.J.P., F.H., A.H.B., E.H. and M.P.Czech designed the research. A.G. and M.P.Czech wrote the manuscript. A.G., D.J.P., F.H., A.H.B., E.H., M.K., K.R., and D.A.M. performed research. K.R. and D.A.M. designed and performed experiments to measure the sensory and sympathetic nerve activities in control and iAdFASNKO mice.

## CONFLICT OF INTEREST

The authors declare no conflict of interest

---

**Supplementary Figure 1: DNL in white and brown adipocytes is not required to maintain euthermia. (A)** Total body weight and BAT (**B**) and iWAT (**C**) depot weight after 1 week at 6°C. **(D)** Core temperature in acute cold challenge (6°C), starting from room temperature in control and iAdFASNKO mice. Graphs show the mean +/− SEM. N= 5-7 mice per group.

**Supplementary Table 1: Primer sequences used in qRT-PCR analysis**

